## Supplementary Figure 1 for "*Atm* loss does not radiosensitize a primary mouse model of *Pten*-deleted brainstem glioma"

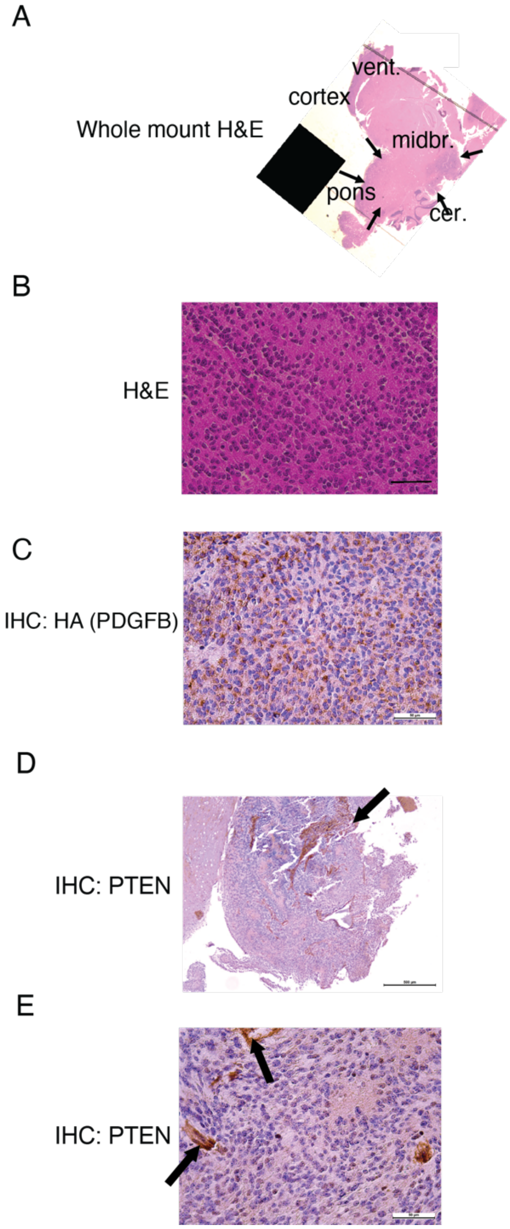


**Supplementary Figure 1. Characterization of nPtenA$^{FL/+}$ brainstem gliomas.** (A) Whole-mount H&E slide showing expansile tumors in the brainstem of nPtenA$^{FL/+}$ mice. Midbr., midbrain; vent, lateral ventricle; cer., cerebellum.

(B) Magnified H&E slides for tumors from nPtenA$^{FL/+}$ mice. Scale bar represents 50 $\mu$m.

(C) Immunohistochemistry staining for HA-tagged PDGFB in tumors from nPtenA$^{FL/+}$ mice. Scale bar represents 50 $\mu$m.

(D) 5X mount showing immunohistochemistry for PTEN of tumors centered in the brainstem (pons) with lack of PTEN reactivity, and diffuse infiltrating borders (arrows) for tumors from nPtenA$^{FL/+}$ mice. Scale bar represents 500 $\mu$m.

(E) 40X mount shows complete loss of PTEN in tumor cells but not normal blood vessels (arrows) in nPtenA$^{FL/+}$ mice. Scale bar represents 50 $\mu$m.
